# Historical biogeography of *Cannabis* in the Iberian Peninsula: palynological evidence

**DOI:** 10.1101/2022.09.16.508344

**Authors:** V. Rull, F. Burjachs, J.S. Carrión, A. Ejarque, S. Fernández, J.A. López-Sáez, R. Luelmo-Lautenschlaeger, J. Ochando, S. Pérez-Díaz, J. Revelles, S. Riera, S. Rodríguez

## Abstract

The tempo and mode of colonization of the Iberian Peninsula (IP) by *Cannabis sativa*, its further internal spreading and the potential cultural and environmental factors involved remain unknown. The available continental-wide European meta-analyses using pollen and archaeological evidence account for only a few IP sites, insufficient for a sound assessment. This paper presents a nearly comprehensive database of almost 60 IP sites with palynological evidence of *Cannabis* and analyzes the corresponding spatiotemporal patterns. The first scattered records of this pollen type, likely corresponding to wild *Cannabis*, date from the Middle and Upper Paleolithic (150 to 12 ky BP) and would have entered the IP by maritime Mediterranean or terrestrial continental pathways, or both. A first burst of introductions, probably in a cultivated form, would have occurred during the Neolithic (7-5 ky BP) using similar paths. Human participation in this Neolithic acceleration remains unclear but cannot be dismissed. A period of reduced *Cannabis* arrivals (mostly via MP) occurred between the Chalcolithic and the Roman Epoch (4.5-2 ky BP), when the innermost parts of the IP were colonized (Late Bronze). A second, likely anthropogenic, introduction acceleration took place in the Middle Ages (1.5 ky BP onward) using the MP and CP. Maximum cultivation and hemp retting activity was recorded during the Modern Ages (16^th^-19^th^ centuries), coinciding with the increased demand of hemp fiber to supply the Spanish royal navy for imperial expansion and commerce. A potential link between *Cannabis* colonization/introduction bursts and climatic warmings has been observed that should be tested with future studies. Regional moisture variations seem to be less influential. Further efforts to enhance and improve the database used in this study are encouraged. The results of this paper should be compared with archaeological and historical evidence to clarify the role of human migrations and cultural changes in the historical biogeography of *Cannabis* in the IP.

## 1. Introduction

*Cannabis*, currently represented by the extant species *C. sativa*, is one of the most ancient crops that has been an integral part of human life since its domestication and continues into the present (Duvall, 2014; Warf, 2014; Small, 2015). Intensive artificial (human-mediated) selection of this plant has created a wide array of varieties and biotypes for diverse uses, including hemp fiber (cordage, textiles, paper, building materials), human and animal feeding (seeds, oil), biofuel, medicine (herbal remedies, pharmaceuticals) recreational drugs (marihuana, hashish), and ritual activities (healing, life cycle rituals, inebriation), among others (Clarke & Merlin, 2013, 2016). Evolutionarily, *Cannabis* is thought to have originated after divergence from its sister genus *Humulus* (both belonging to the family Cannabaceae) during the Oligo-Miocene (28-21 Ma) in the NE Tibetan Plateau and differentiated into their extant subspecies (*C. sativa* subsp. *sativa* and *C. sativa* subsp. *indica*) in the Middle Pleistocene, approximately 1 Ma ago (McPartland, 2018; McPartland et al., 2019). Human domestication of *Cannabis* would have occurred in the early Neolithic (12 kyr BP) of Eastern Asia, which places this plant among the oldest human domesticates (Ren et al., 2021). The human diffusion of *Cannabis* occurred in postglacial times, with a first phase (10-2 kyr BP) of Eurasian dispersal, a further spread into Africa and SE Asia (2-0.5 yr BP) and a diffusion into the Americas from Europe and Asia (1545-1945 CE) (Clarke & Merlin, 2013).

The fossil record of *Cannabis* is largely based on pollen, which is the most extensively used evidence to reconstruct the historical biogeography of this plant and to trace its cultural relationships. However, the morphological similarity between *Cannabis* and *Humulus* pollen has been a handicap for the correct identification of these genera in past records, which is a critical point as the species of these genera have very different ecological requirements and cultural connotations (Portillo et al., 2020; Rull, 2022). This is why different authors refer to the fossil/subfossil representatives of this pollen type using unspecific names such as *Cannabis/Humulus* (thereafter *C/H), Cannabis-type, Humulus-type* or Cannabaceae, among others (McPartland et al., 2018). Several morphological traits have been proposed to be useful to distinguish *Cannabis* and *Humulus* pollen (e.g., Godwin, 1967; Whittington & Gordon, 1987; Whittington & Edwards, 1989; Dörfler, 1990; Fleming & Clarke, 1998; Mercuri et al., 2002; Rull & Vegas-Vilarrúbia, 2014; Trapote et al., 2018) but there is no a universal agreement in this respect and different authors use different criteria.

McPartland et al. (2018) proposed another approach −also called the assemblage approach (Rull, 2022) – according to which *C/H* pollen found in pre-Neolithic steppe assemblages dominated by grasses, chenopods and *Artemisia* should be identified as pollen from wild *Cannabis*, whereas the same morphological pollen type found in association with elements from deciduous mesophytic forests, such as alders (*Alnus*), willows (*Salix*) and poplars (*Populus*), should correspond to wild *Humulus* pollen. According to the same authors, *C/H* pollen from Neolithic and post-Neolithic assemblages dominated by cereals (*Avena, Hordeum, Secale, Triticum*) and/or other anthropic indicators (notably *Centaurea, Polygonum* and *Plantago*) should be associated to cultivated *Cannabis*, especially when *C/H* pollen appears de novo or increases twofold over earlier records.

Using pollen evidence, McPartland et al. (2018) proposed that *Cannabis* could have reached Europe in its wild form and, hence, it could have also been domesticated in this continent. The argument is essentially chronological and is based on the occurrence of *Cannabis* pollen in Europe during Middle-Late Pleistocene (420-40 kyr BP) (Fig. 1). According to these authors, European domestication could have occurred in present-day Bulgaria (eastern Europe) during the late Neolithic-Copper age (7-5 kyr BP) and the expansion across Europe during Bronze-Iron ages (4.5-2.3 kyr BP). This was consistent with the view of Clarke & Merlin (2013), who proposed that the precursor of European hemp (related to *Cannabis sativa* subsp. *sativa*) would have been centered in the Caucasus region, from where it would have dispersed westward during the Pleistocene. This would also be chronologically consistent with the idea of Pleistocene (^~^1 Ma) divergence of *C. sativa* subsp. *sativa* and *C. sativa* subsp. *indica* (McPartland, 2018) and the further expansion of the first into Europe. All these considerations about the possibility of European domestication were based on the reconsideration of palynological evidence by McPartland et al. (2018), using the assemblage approach for the attribution of *C/H* pollen in the available studies to either one or another of these genera, in their wild and/or cultivated forms. Archaeological evidence at continental level was also consistent with this reinterpretation (McPartland & Hegman, 2018).

**Figure 1.**
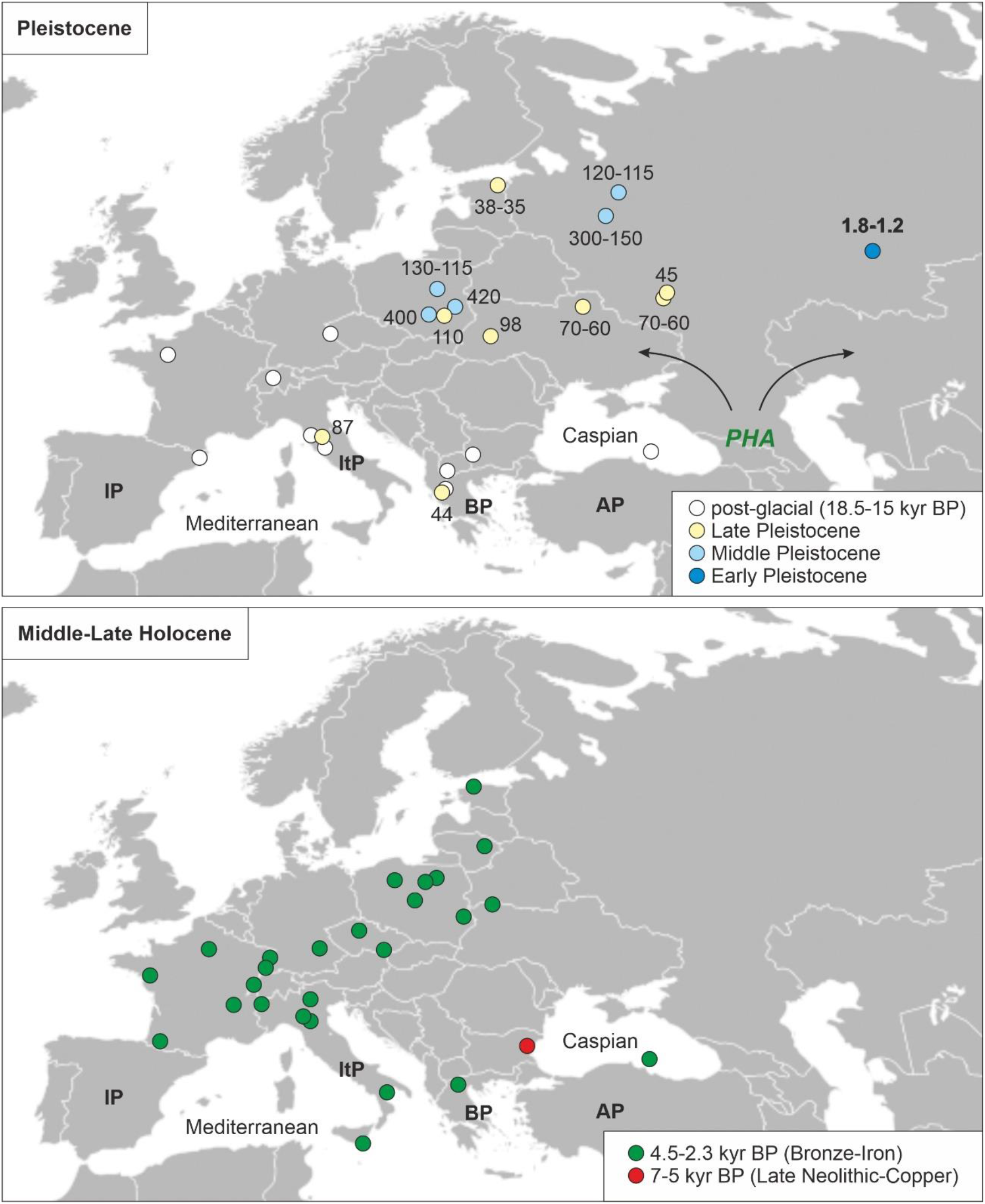
Pleistocene and Holocene European pollen records of *Cannabis*, identified using the assemblage approach. PHA is the precursor of the European *Cannabis* according to Clarke & Merlin (2013). Ages in million years before present (bold numbers and in thousand years before present (normal numbers). Redrawn from Rull (2022), based on raw data from McPartland et al. (2018). AP, Anatolian Peninsula; BP, Balkan Peninsula; IP, Iberian Peninsula; ItP, Italian Peninsula.

The same palynological meta-analysis showed that *Cannabis* pollen was already present in western Europe, including the NE Iberian Peninsula (thereafter IP), during post-glacial times (18.5-15 kyr BP) (Fig. 1), indicating that Pleistocene *Cannabis* dispersal would have occurred prior to its eastern domestication. This implies that at least two westward dispersal events of *Cannabis* could have occurred in Europe, one in wild state (18.5-15 kyr BP) and another in its cultivated form (4.5 kyr BP onward). In spite of its biogeographical and cultural relevance, the Iberian Peninsula is not well represented in analyses and meta-analyses about *Cannabis* history (Rull, 2022). For example, Clarke & Merlin (2013) used a single site (Lake Estanya) to infer the arrival of *Cannabis* into the IP by 600 CE, which strongly contrasts with the records of McPartland et al. (2018), who used 10 IP sites from peer-reviewed papers and the European pollen database, and identified post-glacial (18.5-14.5 kyr BP) *Cannabis* pollen in two of them (Lakes Banyoles and La Roya) and Holocene pollen of this type in the other nine, ranging from 10 to 0.8 kyr BP. According to Rull (2022), more sites are available in the IP for developing analyses of this type that remain unnoticed in the available investigations and should be addressed for a better appraisal.

This paper uses a newly obtained and more complete palynological dataset on the spatiotemporal patterns of occurrence and abundance of *Cannabis* pollen across the IP. The raw data has been retrieved after an exhaustive literature review including not only records from international publications and continental-wide databases, but also results published in local/regional journals and dissertations, as well as unpublished data from the author’s own data files (original counts and spreadsheets) that had not been reported in previous publications due to the scarcity of *C/H* pollen and/or the lack of interest on this pollen type at the time of publication. The final dataset contains nearly 60 sites. In this paper, this dataset is analyzed with focus on the following research questions about *Cannabis* in the IP: (i) the arrival timing, (ii) the geographical origin and the penetration pathways, (iii) the wild or domesticated nature of the incomers, (iv) the further expansion across the IP, (v) the potential relationships with climatic changes, and (vi) the onset and development of large-scale cultivation and retting. Inferences about cultural relationships need archaeological and historical evidence to be properly understood and will be addressed in future studies. This paper aims to provide the palynological basis for these integrated studies.

## 2. The Iberian Peninsula: biogeographical and historical context

The IP is a key biogeographical region due to its high biodiversity and endemism levels, along with its transitional character between contrasting biogeographical regions, which results in a very peculiar biota in the European context. The IP is one of the main centers of Mediterranean plant diversity, together with the Anatolian, the Balkan and the Italian peninsulas (Médail and Quezel, 1997) (Fig. 1). These peninsulas have had a fundamental role not only as biodiversity cradles but also as glacial refugia, especially during the Last Glacial Maximum (LGM) (Hewitt, 1999; Birks, 2016). It has been estimated that roughly a quarter of the LGM refugia have been located in the IP and the Balearic Islands (Médail & Diadema, 2009). In this context, the Mediterranean Region, and in particular the IP (González-Sampériz et al., 2010; Carrión et al., 2003, 2012, 2022), emerge as a singular territory in terms of geographical isolation and persistence of populations and species.

The present Iberian vascular flora is especially rich – almost 6300 species (^~^740 non-native), distributed in nearly 1300 genera and 190 families – and shows a high degree of specific endemism (^~^25% of the native flora). This represents more than 50% of the European flora. The main Iberian centers of diversity and endemism are situated in the mountain ranges and the richest groups are the Asteraceae, the Fabaceae and the Poaceae, whereas the Gimnosperms and the Pteridophytes are the less represented (Aedo et al., 2017). Traditionally, the IP has been subdivided into two major biogeographical domains characterized by contrasting bioclimatic features: the Mediterranean and the Eurosiberian regions. The Mediterranean region is characterized by the occurrence of Mediterranean macrobioclimates, with at least two consecutive arid months during the warmest season (summer), and dominates most of the IP (Fig. 2). The Eurosiberian region is restricted to the N and NW sectors of the peninsula (^~^20% of the total surface) and is characterized by wet temperate macrobioclimates, without a period of two or more consecutive with summer aridity (Rivas-Martínez et al., 2017). The Mediterranean region is dominated by species distributed across southern Europe and northern Africa, whereas the Eurosiberian region is characterized by species distributed across central and northern Europe (Aedo et al., 2017).

**Figure 2.**
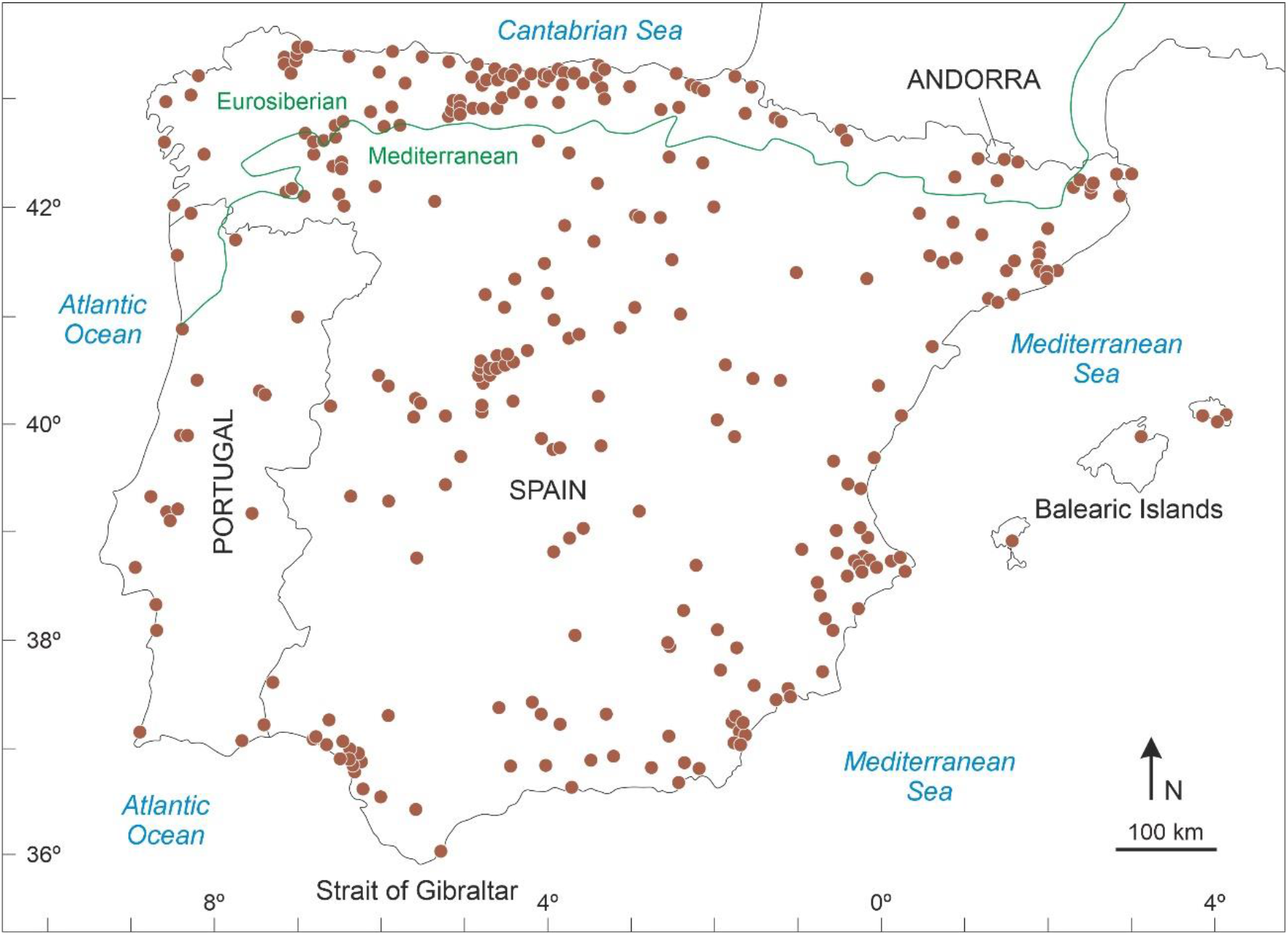
Map showing the sites with Pliocene to Quaternary pollen records from the Iberian Peninsula, according to Carrión et al. (2012). Marine cores have not been included. The approximate boundary between the Eurosiberian and Mediterranean biogeographical regions is indicated by a green line.

Land use is also strongly influenced by bioclimatic and biogeographical patterns. In the Eurosiberian region, land use is comparable to the rest of Atlantic Europe, where the traditional agrarian economy is based on cattle raising for milk and meat, combined with small-scale crops such as maize, potatoes and other vegetables, as well as fruits such as apples and chestnuts. The current landscape is a mosaic of small fields, woodlands and pastures, combined with larger areas of heathlands, forests and grasslands, especially in higher elevations. The Mediterranean Iberia is dominated by a totally different landscape resulting from land-use practices adapted to the occurrence of the climatic summer drought. In these conditions, crops not needing irrigation are favored. Annual crops, especially cereals, largely dominate the central IP, which is characterized by huge medium-elevation flat terrains. In the southern half, where climates are not so cold, typical Mediterranean crops such as vineyards and olive grows dominate the landscape. In the Mediterranean region, extensive crops are the norm and animal husbandry occupies a subordinate position, except in some low-mountain areas, where extensive parkland landscapes dominate (Loidi, 2017). The Mediterranean biome and its climatic features are better suited than its Eurosiberian counterpart for the growth of *Cannabis*.

The oldest human remains found in the Iberian Peninsula date from approximately 1.4-1.2 Ma (Early Pleistocene) and correspond to *Homo antecessor*, an autochthonous species related to *H. erectus* (Bermúdez de Castro et al., 1997), and an undetermined hominin specimen known as BL02-J54-100 (Toro-Moyano et al., 2013). Heidelbergs and their relatives Neanderthals entered the IP at least 0.6 Ma (Early Paleolithic) (Carrión et al., 2011; Rosas et al., 2019), and modern humans did the same by 45-30 kyr BP (Carrión et al., 2019). The Neolithization of the IP began at approximately 5800 BCE (^~^7.8 kyr BP), when the former hunter-gatherer nomadic societies were replaced by more stable agricultural-based cultures (García-Martínez de Lagrán, 2015). During the Bronze Age (2700-800 BCE) and the Iron Age (800-70 BCE), a diversity of local cultures developed on the IP, which were influenced by Indo-European migrations, from the north (1^st^ millennium BCE) and the Phoenicians, arriving by the Mediterranean coasts during the 11^th^ century BCE. The autochthonous Iberian culture fully developed during the 7^th^-6^th^ centuries BCE but colonizations and invasions from the Mediterranean area continued with the Greeks (8^th^ century BCE), Cartaginians (5^th^ century BCE) and the Romans, who arrived by 220 BCE and occupied the IP for almost 600 years. After a transitional phase in which the IP was occupied by the Germanic Visigoths, coming from the north, the next long-lasting invasion was that of the Muslims, who entered the IP from Africa and remained in the IP for almost eight centuries (8^th^-15^th^ centuries CE). The Late Antiquity and the Middle Ages (5^th^-15^th^ centuries CE) signified the progressive southward expansion of Christian cultures and the unification of monarchies, followed by the consolidation of the modern geographical and political configuration. After the Middle Ages, the kingdoms of the IP shifted from colonized to colonizer countries and expanded their dominions all over the world, with emphasis on the American continent, a situation that remained until the 19^th^-20^th^ centuries CE (Marín, 1995).

This brief summary shows that the IP is a strategic spot in biogeographical, cultural and historical terms, mainly due to its geographic location, which has promoted the interaction of a diversity of cultures over prehistoric and historic times, favored by numerous and varied land and sea connections. Therefore, it might be expected that the IP would also be a relevant place for the study of a plant such as *Cannabis*, which has been subjected to intensive and extensive natural and human-mediated dissemination.

## 3. Methods

The newly obtained database for the presence and abundance of *C/H* pollen in the IP, in its first version called CHIP 1.0, consists of 56 sites situated in the IP and the Balearic Islands, including localities from Spain, Portugal and Andorra. The initial basis for the assembling of this database was the first version of the Iberian Paleoflora and Paleovegetation (Carrión et al., 2012), whose most recent updating is in progress (Carrión et al., 2022) and has also been used to improve the CHIP database. The Iberian Paleoflora and Paleovegetation database gathers most pollen records available for the IP, encompassing ^~^330 sites in the first version and ^~^420 sites in the second version (Fig. 2). This database was used to locate the sites with published and unpublished *C/H* pollen records. The published records were extracted directly from the original publications, whereas the unpublished results were obtained from the personal records of the involved researchers (counting sheets, spreadsheets). Not all researchers could be contacted and, therefore, it cannot be ruled out that some information is missed. However, to the knowledge of the authors of this paper, all the relevant data have been included here and eventual omissions are considered to be minor and not affecting the general conclusions. In spite of this, the authors of this paper remain alert to possible additions for the improvement and updating of the CHIP database.

As usual, the *C/H* pollen has been labelled using a variety of unspecific names by different authors and have used the generic name *Cannabis* only in a few cases (*C. sativa* in a single case) with no mention of the criteria used, with a few exceptions (e.g., Rull & Vegas-Vilarrúbia, 2014; Servera-Vives et al., 2018; Trapote et al., 2018). Due to the general unavailability of the original material (pollen slides, residues) to reexamine these identifications, we considered the assemblage approach of McPartland et al. (2018) and the pollen of each record tried to be assigned to wild *Cannabis* (pre-Neolithic open vegetation, especially steppes), wild *Humulus* (pre-Neolithic woodlands, especially deciduous forests) or cultivated *Cannabis* (Neolithic and post-Neolithic croplands and anthropized open vegetation), according to the original interpretations. This approach could not be applied in many cases, especially in sites with single and occasional occurrences, which hindered the use of quantitative criteria (i.e., the twofold increase with respect to pre-anthropogenic values). In spite of this, the assemblage approach was useful in selected cases, which are discussed on an individual basis.

## 4. Results

The final dataset used in this study is summarized in Table 1 and is provided in full as a spreadsheet in the Supplementary Material. Fig. 3 shows a clear NW-SW gradient with a higher density of sites in the NE sector (including the Balearic Islands) and an outstanding absence of *C/H* records in the southern area, with the exception of coastal environments. The occurrence of the interior SW “blind” *C/H* area cannot be attributed to the absence of palynological studies (Fig. 2) but likely to the actual lack of *C/H* records. Whether this is due to the absence of this pollen type or to its deliberate/unintentional oversight remains unknown, although the second possibility would be inconsistent with the usual palynological procedures in the IP (Carrión et al., 2012).

**Table 1.** Data used for the spatiotemporal analysis of *Cannabis/Humulus* pollen across the Iberian Peninsula, according to the original references (see Fig. 3 for the location of sites and the Supplementary Material for more details). Elev (m), elevation in m a.s.l.; OP, occurrence pattern (C, continuous; D, discontinuous; O, occasional; S, single occurrence); MA, maximum abundance (+, <1%); FA, first appearance. ND, no data.

| Site (code) | Lat N | Long | OP | MA (%) | FA | Period | References |
| --- | --- | --- | --- | --- | --- | --- | --- |
| Addaia (AD) | 39°59'23" | 4°12'22"E | O | + | 2500 BCE | Chalcolithic | Servera-Vives et al. (2018) |
| Alcúdia (AL) | 39°47'48" | 03°06'24"E | D | + | 6270 <sup>14</sup>C BP | Neolithic | Burjachs et al. (1994, 2017) |
| Algendar (AG) | 39°56'26" | 03°57'31"E | C | + | 9000 cal BP | Mesolithic | Burjachs et al. (2017) |
| Antas (AN)* | 37°12'30" | 01°49'25"W | S | ND | ND | Mesolithic | Pantaleón-Cano et al. (2003) |
| Asperillo (AS) | 37°02'34" | 06°38'04"W | S | + | 24,800-23,200 cal BP | Upper Paleolithic | Fernández et al. (2021) |
| As Pontes (AP) | 43°21' | 07°29'W | S | 10 | 3500 BCE | Neolithic | López-Sáez et al. (2003a, b) |
| Aulaga (AU) | 36°59'51" | 06°25'49"W | O | + | 8000-7000 cal BP | Mesolithic | Manzano et al. (2018) |
| Banyoles 1 (BA1) | 42°07'34" | 02°45'32"E | D | + | 31,000 cal BP | Upper Paleolithic | Pérez-Obiol & Julià (1994) |
| Banyoles 2 (BA2) | 42°07'34" | 02°45'32"E | S | + | ND | Neolithic | Revelles et al. (2016) |
| Banyoles 3 (BA3)* | 42°07'45" | 02°45'07"E | D | + | 8300 cal BP | Mesolithic | Revelles et al. (2015) |
| Besós (BE) | 41°24' | 02°15'E | C | + | 500 CE | Medieval | Riera & Palet (2005, 2008) |
| Bolomor (BO) | 39°03' | 00°15'W | O | + | 150 ky BP | Middle Paleolithic | Ochando et al. (2019) |
| Cañamares (CA) | 41°11'59" | 02°57'25"W | S | + | 1450 CE | Medieval | Currás (2012) |
| Carcavai (CR) | 37°03'35" | 08°04'48"W | O | + | 4500 cal BP | Chalcolithic | Schneider et al. (2010, 2016) |
| Castelló (CS) | 42°16'48" | 03°06'09"E | C | + | 500 CE | Medieval | Ejarque et al. (2016) |
| Creixell (CX) | 41°09'41" | 01°27'16"E | D | + | 5000 cal BP | Neolithic | Burjachs & Schulte (2003), Burjachs (2012) |
| Cubelles (CU) | 41°13' | 01°39'E | D | + | 5000 <sup>14</sup>C BP | Neolithic | Riera & Esteban (1997), Burjachs (2012) |
| Cueto Espina (CE)* | 43°04'01" | 03°55'06"W | D | 5 | 5720 cal BP | Neolithic | Rodríguez-Coterón (PhD, unpublished) |
| Eivissa (EI) | 38°55'01" | 01°27'08"E | S | + | 650 cal BP | Medieval | Yll et al. (2009), Carrión et al. (2012) |
| Elx (EX)* | 38°10'28" | 00°45'10"W | D | + | 9420 cal BP | Mesolithic | Carrión et al. (2012) |
| Empúries (EM) | 42°16'53" | 03°07'13"E | D | + | 3800 cal BP | Chalcolithic | Burjachs et al. (2005), Burjachs (2012) |
| Estanya 1 (ET1) | 42°01'44" | 00°31'44"E | C | 25 | 600 CE | Medieval | Riera et al. (2004, 2006) |
| Estanya 2 (ET2) | 42°01'44" | 00°31'44"E | C | 5 | 1000 cal BP | Medieval | Morellón et al. (2011), González-Sampériz et al. (2017) |
| Estanyons (EN) | 42°28'50" | 01°37'45"E | O | + | 7000 cal BP | Neolithic | Miras et al. (2007), Ejarque et al. (2010) |
| Fronteras (FR) | 40°14'15" | 03°41'48"E | S | + | ND | ND | Macías et al. (1996) |
| Gallocanta (GA) | 40°58'25" | 01°30'22"W | D | + | 1110 CE | Medieval | Burjachs et al. (1996) |
| Gesaleta (GE)* | 42°58'54" | 01°34'17"W | O | + | 11,420-10,940 cal BP | Upper Paleolithic | Ruiz-Alonso et al. (2019) |
| Gran Dolina (GD) | 42°21'07" | 03°31'12"W | S | + | ND | Middle Pleistocene | García-Antón (1989) |
| Grau (GR)* | 39°56'53" | 04°15'31"E | S | + | 1970 cal BP | Roman | Carrión et al. (2012), Burjachs et al. (2017) |
| Gusó (GU) | 40°05'31" | 3° 05' 23" E | O | + | 4000 <sup>14</sup> C BP | Chalcolithic | Blech et al. (1998) |
| La Cruz (LC) | 39°59'16" | 01°52'25"W | D | + | >1350 CE | Medieval | Burjachs (1996), Julià et al. (1998) |
| La Roya (LR) | 42°13'00" | 06°46'00" W | D | + | 14,500-10,500 cal BP | Upper Paleolithic | Allen et al. (1996), Muñoz-Sobrinho et al. (2013, 2018) |
| Mirador (MI) | 42°20'58" | 03°30'33" W | S | + | 5900-5700 cal BP | Neolithic | Expósito (2006) |
| Montcortès (MO) | 42°19'50" | 00°59'41" E | C | 60 | 600 CE | Medieval | Rull et al. (2021a, b), Trapote et al. (2018) |
| Orris Setut (OS) | 42°28'57" | 01°39'01" E | ND | + | 1500 CE | Modern Age | Ejarque et al. (2010) |
| Perales (PS) | 40°19'25" | 03°39'26" E | O | + | ND | Late Bronze Age | Macías et al. (1996) |
| Pocito Chico (PC) | 36°40'43" | 06°14'51"W | S | 5 | ND | Neolithic | López-Sáez et al. (2001, 2002) |
| Port Lligat 1 (PL1) | 42°17'30" | 03°17'24"E | D | 5 | 1530 CE | Modern Age | López-Sáez et al. (2009) |
| Port Lligat 2 (PL2) | 42°17'37" | 03°17'25"E | C | + | 4800 cal BP | Chalcolithic | López-Merino et al. (2017) |
| Porter (PT)* | 39°52'14" | 03°17'25"E | D | + | 5570-5400 cal BP | Neolithic | Yll et al. (1997) |
| Pradell (PR) | 42°17'20" | 04°07'53"E | C | + | 1500 CE | Medieval | Ejarque et al. (2009) |
| Quarteira (QU) | 37°06'00" | 08°08'15"W | O | + | 4500 cal BP | Chalcolithic | Schneider et al. (2010, 2016) |
| Rascafría (RA) | 40°54'42" | 03°51'47"W | S | + | 1500 cal BP | Medieval | Franco-Múgica et al. (1988) |
| Reixach (RE) | 42°00'46" | 03°05'03"E | C | + | 5600 <sup>14</sup>C BP | Neolithic | Blech et al. (1998), Burjachs (2012) |
| Riu Orris (RO) | 42°29'20" | 01°38'14"E | C | + | 600 CE | Medieval | Ejarque et al. (2010) |
| Roquetas (RQ)* | 36°47'40" | 02°35'20"W | S | + | 2400 cal BP | Iron Age | Pantaleón-Cano et al. (2003) |
| San Rafael (SR)* | 36°46'25" | 02°36'05"W | S | + | 6360 cal BP | Neolithic | Pantaleón-Cano et al. (2003) |
| Sanabria (SA) | 42°07'03" | 06°43'00"W | O | + | <2400 cal BP | Iron Age | Julià et al. (2007) |
| Sils (SI) | 41°48'00" | 02°44'00"W | D | 5 | ND | ND | Pérez-Obiol et al. (1991) |
| Sobrestany (SO) | 42°05'27" | 03°07'22"E | D | 5 | 6300 cal BP | Neolithic | Parra et al. (2005), Burjachs (2012) |
| Somolinos 1 (SM1) | 41°15'05" | 03°03'54"W | O | + | 1700 CE | Modern Age | Currás (2012) |
| Somolinos 2 (SM2) | 41°14'48" | 03°03'43"W | O | + | 600 BCE | Iron Age | Currás (2012) |
| Son Bou (SB)* | 39°55'29" | 04°01'28"W | D | + | 13,400 cal BP | Upper Paleolithic | Yll et al. (1997) |
| Teixoneres (TE) | 41°48'25" | 02°09'02"E | O | + | 100 ky BP | Middle Paleolithic | Ochando et al. (2020a) |
| Toll (TO) | 41°48'25" | 02°09'02"E | O | + | 43 cal ky BP | Upper Paleolithic | Ochando et al. (2020b) |
| Veranes (VE) | 43°29'03" | 05°45'13"W | O | + | 1<sup>st</sup>-4<sup>th</sup> centuries | Roman | López-Sáez (2012) |

\*Unpublished data, retrieved from author's personal files

**Figure 3.**
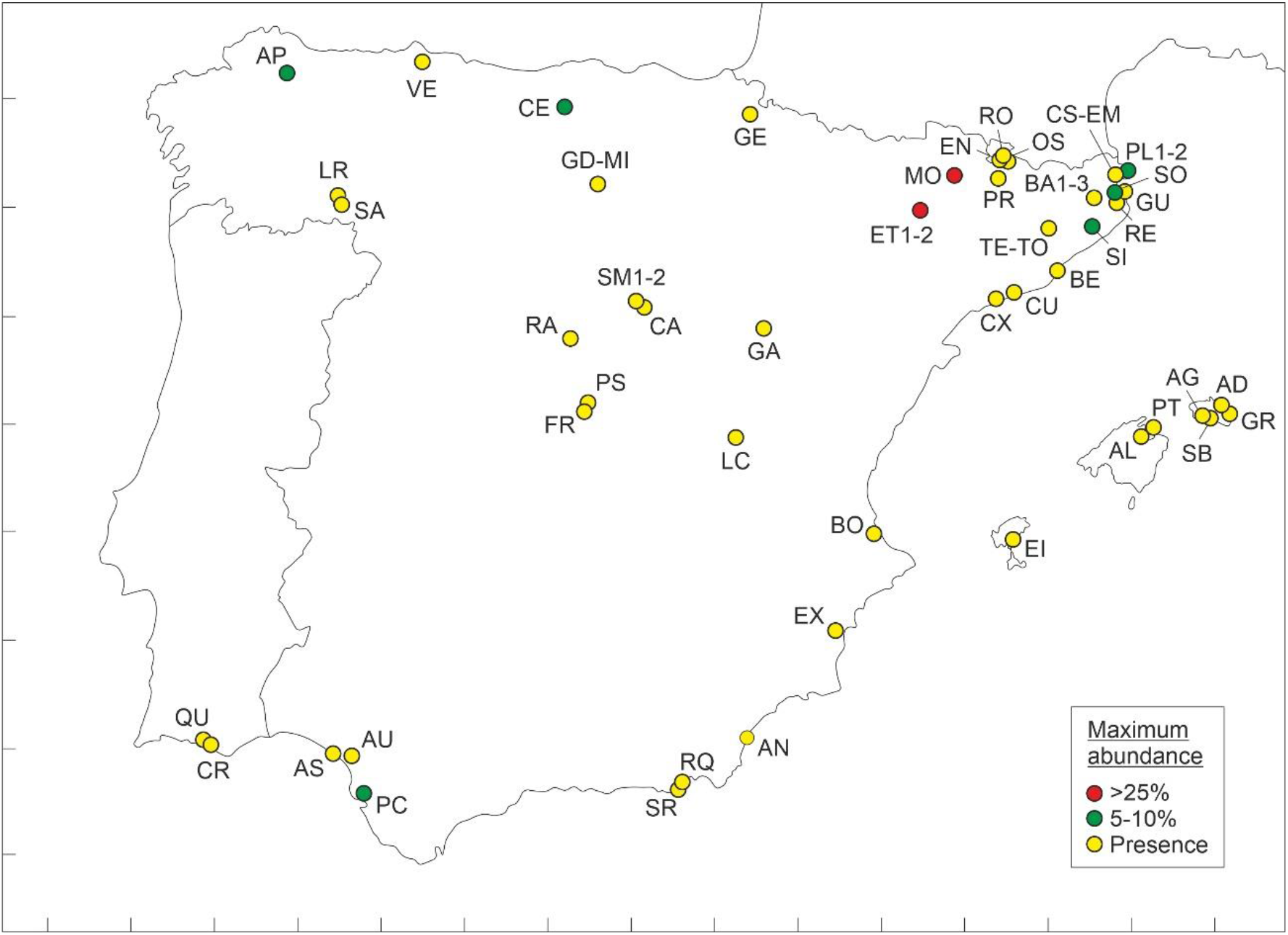
Map of the Iberian Peninsula showing the sites used in this study, with indication of the maximum abundances of *C/H* pollen recorded. See the Supplementary Material for more details.

### 4.1. Abundance and continuity

Overall, continuous *C/H* records occurred in 10 cases, of which only two (Estanya and Montcortès) showed values over 1%. These two sites hold the highest *C/H* pollen values with maxima between ^~^25% and ^~^60%, respectively. Discontinuous occurrences were more frequent (16), with four sites (Cueto Espina, Port Lligat, Sils and Sobrestany) attaining maxima of 5% or above. Occasional and single occurrences were the most frequent (29 sites), with only two sites with maximum abundances of 5% (As Pontes, Pocito Chico) and the remaining below the “presence” boundary (1%). Therefore, the most frequent case was that of single/occasional occurrences documenting only presence, which accounted for roughly half of the sites (27). Table 2 summarizes the localities with higher abundances of *C/H* pollen and their main features. Noteworthy, all of them lie in the northernmost part of the IP except one (Pocito Chico), which is the south. From a chronological perspective, *C/H* pollen was present in most of these localities since the Neolithic, except Estanya and Montcortès, where this pollen appeared in the Middle Ages. Using the assemblage approach of McPartland et al. (2018), *C/H* pollen from sites with higher abundances was assigned to cultivated *Cannabis*, except two (Cueto Espina and Sils), where the available information was insufficient to apply this method.

**Table 2.**
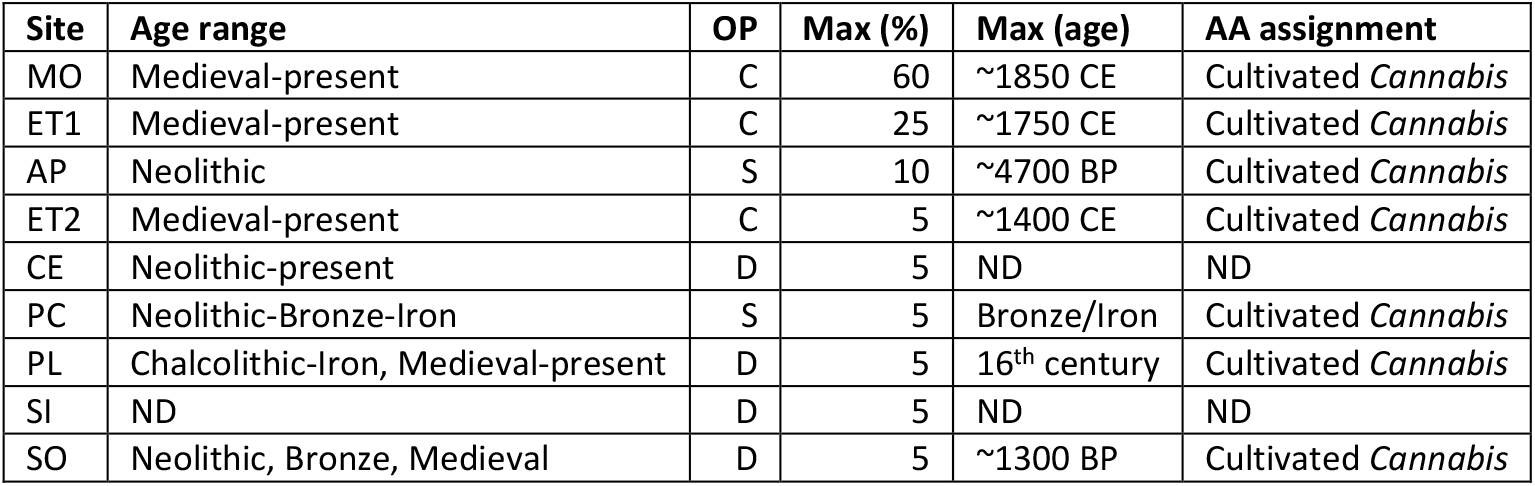
Sites with maximum *C/H* pollen abundances over 5% (see also Fig. 3) and occurrence patterns (OP: C, continuous; D, discontinuous; S, single occurrence). The prehistoric/historical phases of occurrence of *C/H* pollen and the age of the acme in each site is indicated. The taxonomic assignment according to the assemblage approach (AA) of McPartland et al. (2018) is also provided. ND, no data.

| Site | Age range | OP | Max (%) | Max (age) | AA assignment |
| --- | --- | --- | --- | --- | --- |
| MO | Medieval-present | C | 60 | ~1850 CE | Cultivated *Cannabis* |
| ET1 | Medieval-present | C | 25 | ~1750 CE | Cultivated *Cannabis* |
| AP | Neolithic | S | 10 | ~4700 BP | Cultivated *Cannabis* |
| ET2 | Medieval-present | C | 5 | ~1400 CE | Cultivated *Cannabis* |
| CE | Neolithic-present | D | 5 | ND | ND |
| PC | Neolithic-Bronze-Iron | S | 5 | Bronze/Iron | Cultivated *Cannabis* |
| PL | Chalcolithic-Iron, Medieval-present | D | 5 | 16<sup>th</sup> century | Cultivated *Cannabis* |
| SI | ND | D | 5 | ND | ND |
| SO | Neolithic, Bronze, Medieval | D | 5 | ~1300 BP | Cultivated *Cannabis* |

### 4.2. Chronological sequence

Considering the first appearance (FA) of *C/H* pollen, regardless their abundances, the localities follow a stepped trend from the Middle Paleolithic to the Modern Age (Fig. 4), with maxima in the Neolithic and the Middle Ages, when FAs experienced sharp increases (Fig. 5). Before the Neolithic acceleration, *C/H* records are not uncommon (26% of the total), indicating relatively intense colonization of the IP prior to the agricultural revolution, with 17% of the FAs occurring during the Middle and Upper Paleolithic. The period following the Neolithic was characterized by a significant decline of FAs, which attained minimum values between the Bronze Age and the Roman period. The Medieval acceleration was more intense than the Neolithic rise and was followed by a new decline during the Modern Age, when the last FAs were recorded.

**Figure 4.**
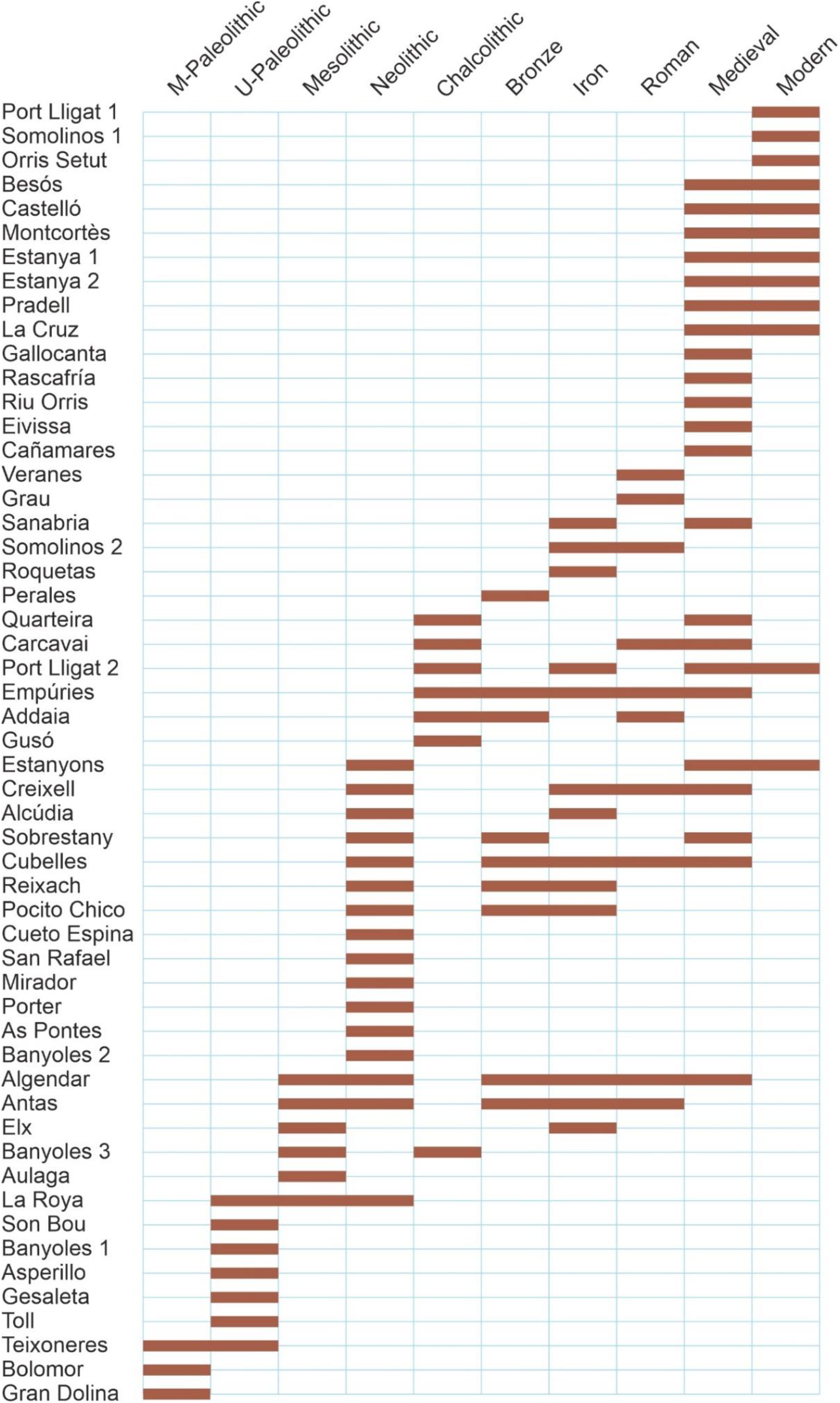
Histogram of the first appearances (FAs) of *C/H* pollen sorted by prehistoric/historical phases.

**Figure 5.**
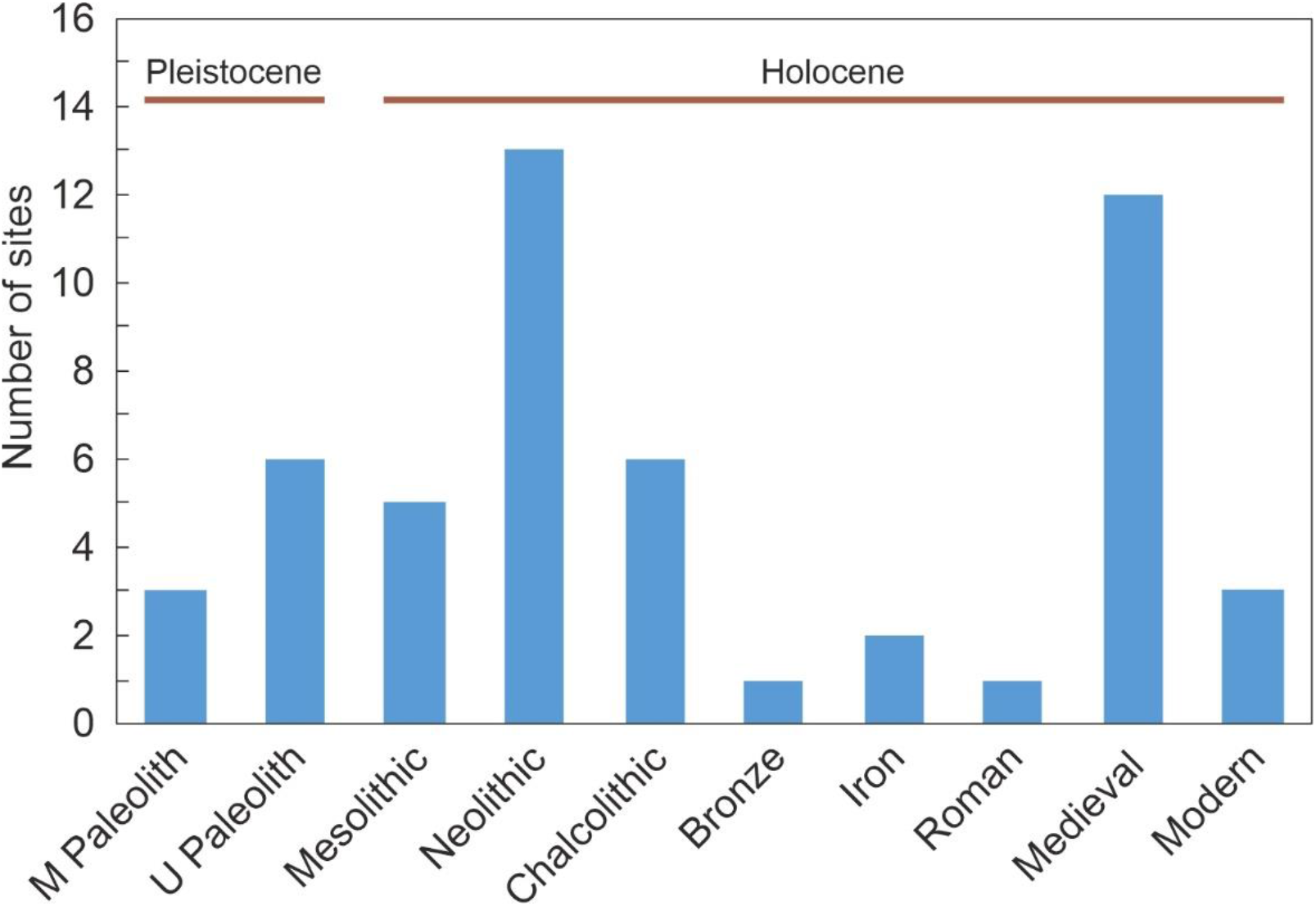
Chronological ranges of *C/H* pollen by site. Fronteras and Sils are not represented due to lack of data.

### 4.3. Spatiotemporal patterns

When the FAs are depicted geographically by cultural phases, some interesting spatiotemporal patterns emerge (Fig. 6). Paleolithic/Pleistocene records are mostly peripheral to the IP, with the older records situated in the Mediterranean coasts, especially in the NE sector, and the younger records in the western IP sector. The lack of *C/H* records in the central part of the peninsula is noteworthy. During the Mesolithic, all FAs were situated along the Mediterranean coasts with no arrivals in the rest of the IP. The Neolithic acceleration was characterized by the concentration of FAs in the northern part of the IP, with the older and most frequent appearances in the NE sector and progressively younger ages toward the NW. Also noteworthy is a relatively old FA recorded in the SE area. The central IP remained uncolonized by *C/H* pollen, even during the Neolithic expansion quickening. The comparatively low Chalcolithic arrivals were concentrated in the NE and SW sector, with nothing in between. During the phase of minimum arrivals (Bronze Age to Roman period; Fig. 5), a more widespread pattern of FA was recorded, with the first appearances of *C/H* pollen in the central IP. Finally, during the Medieval burst, most FAs were situated in a gradient from the northeastern to the central IP, indicating a second major colonization process following this direction.

**Figure 6.**
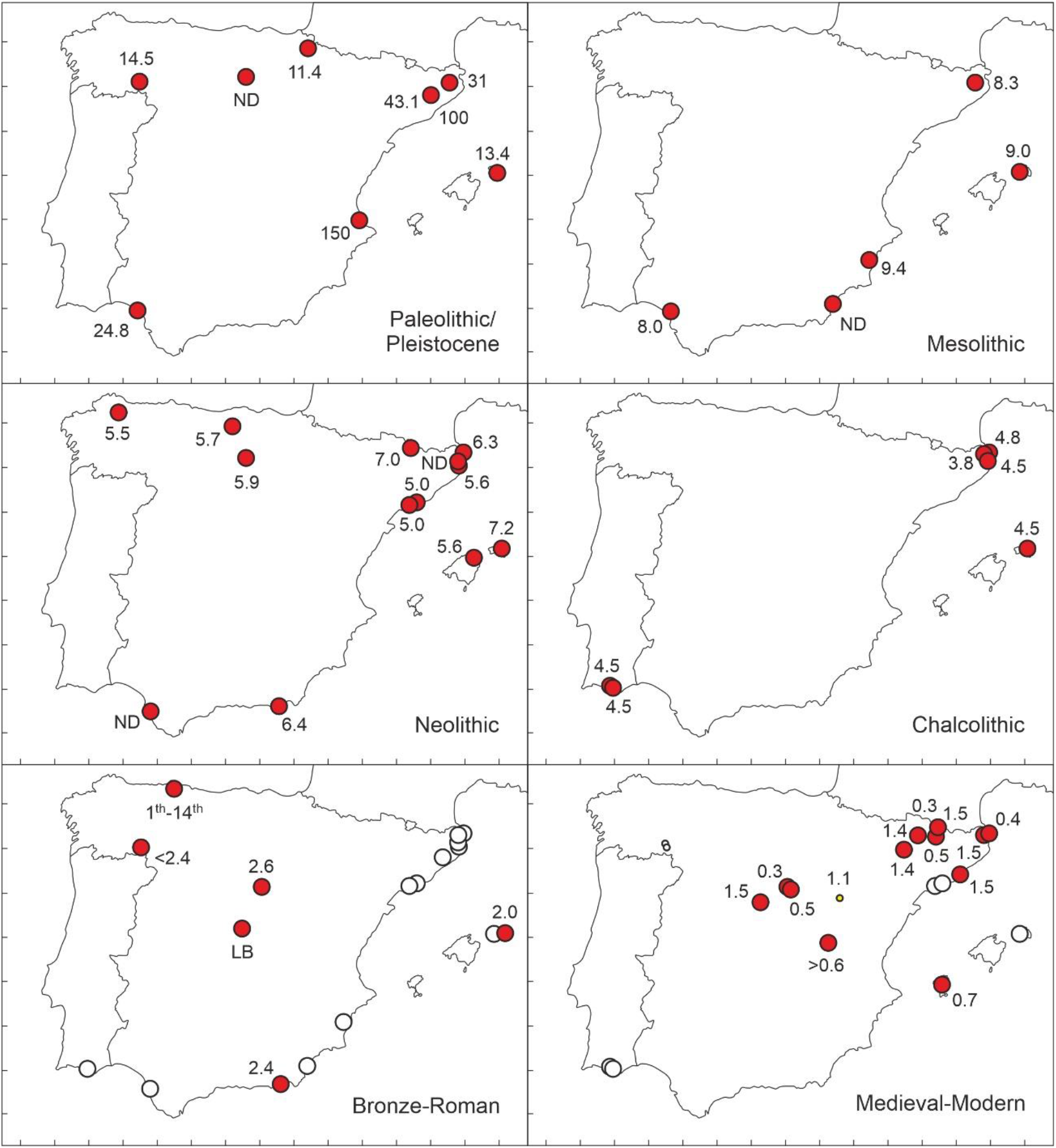
Spatiotemporal patterns of the first appearances (FAs) of *C/H* pollen subdivided into the main cultural phases. Numbers are ages in years BP. Sites with *C/H* pollen continuity after the FA are marked by open dots. LB, Late Bronze Age; ND, no precise dating.

## 5. Discussion and conclusions

Spatiotemporal patterns of FAs documented in this analysis suggest the occurrence of three major colonization waves of the IP by *C/H* pollen: (1) a pre-anthropogenic Middle/Upper Paleolithic phase between ^~^150 kyr BP and ^~^12 kyr BP with a possible east-west component; (2) a Neolithic colonization acceleration (^~^7-5 kyr BP) with a clear NE-NW trend; and (3) a second Medieval burst (1.5 kyr BP onward) proceeding mainly from the NE to the center of the IP. In between, phases of lower colonization intensity but relevant geographical meaning occurred during the Mesolithic, the Chalcolithic, and the Bronze/Iron/Roman phases. All these events and processes will be discussed in more detail in relation to the cultural and environmental history of the IP, but first it is pertinent to assess the palynological evidence required to attest the actual presence of the parent plants of *C/H* pollen and to document the different activities of hemp industry developed in a given locality or region.

### 5.1. Pollen representativeness: local cultivation, hemp retting or long-distance dispersal?

First of all, it should be noted that, as usual in anemophilous pollen types, the presence of *C/H* pollen does not necessarily involve the local presence of the parent plant. Indeed, aerobiological studies have shown, for example, that *Cannabis* pollen from northern African crops can reach the southern IP, situated nearly 100 km north, in significant amounts under favorable meteorological conditions, notably wind direction (Cabezudo et al., 1997; Munuera et al., 2006). Studies on modern pollen sedimentation in one of the localities used in this work (Montcortès) have shown that *Cannabis* pollen abundances of 5-10% can be found in sites with no evidence of local cultivation (Rull et al., 2017). Therefore, low but relevant amounts of *Cannabis* pollen in paleoecological archives from the IP cannot be used to infer the local presence of its parent plants but, rather, their occurrence in the surrounding region. In the present study, based on sedimentary records, most values are below 10% and, therefore, the local occurrence of *Cannabis* pollen sources is not guaranteed. However, in biogeographical studies like this, involving continental-wide movements, the resolution is above 100 km and regional occurrences of this magnitude based on single-site records are representative enough. Similarly, given its strong regional dispersion power, the absence of this pollen in a particular site is considered to be sound evidence for the absence of the parent plants across a ^~^100 km-wide regional range, even under favorable wind conditions. In summary, pollen values of 10% or more are considered to be indicative of local *Cannabis* presence, whereas lower amounts are suggestive of regional presence and, contrary to the well-known assessment that absence of evidence is not evidence of absence, lack of pollen can be considered strong evidence for the absence of *Cannabis* populations in the surrounding region.

More studies are available for pollen percentage as a proxy for hemp retting. The soaking of *Cannabis* plants in lake waters to facilitate the extraction of hemp fibers is an extra source for *C/H* pollen, which is directly incorporated into aquatic sediments and is therefore overrepresented in the resulting fossil assemblages. Values of 80-90% of the pollen sum are not unusual in these cases (Szczepanek, 1971; Peglar, 1993; Hölzer & Hölzer, 1998; Nakagawa et al., 2000; Gearey et al., 2005; Schofield & Waller, 2005; Kittel et al., 2014), but abundances of 15-25% have been considered enough for interpreting hemp retting (Whittington & Gordon, 1987; Peglar, 1993; Mercuri et al., 2002; Riera et al., 2004, 2006; Rull et al., 2011; Demske et al., 2016; Trapote et al., 2018). In this study, only two localities (Estanya and Montcortès) are above this threshold (Table 2). In Montcortès, more detailed studies been performed using genetic evidence for the presence of bacteria actually involved in the fermentation process that detach the hemp fibers from the stalk. These studies have shown that, during the main hemp retting phase extending from 16^th^ to 20^th^ centuries, *Cannabis* pollen abundances were consistently above 20% (Rull et al., 2022). Therefore, this boundary could be tentatively considered a reliable threshold to infer hemp retting in the IP.

Using this evidence, hemp retting can be reliably documented in the Estanya and Montcortès records during Modern and Contemporaneous times (Fig. 7). In both records, *C/H* pollen appeared at the beginning of Middle Ages (^~^600 CE) in abundances that vary between 5 and 10%, which suggests local/regional cultivation (Riera et al., 2004; Rull et al., 2021a, b). In Estanya, a gradual increase by the end of the Middle Ages (^~^1400 CE) culminated in a maximum of 25% during Modern times (^~^1750 CE), which is compatible with hemp retting, followed by a Contemporary drop to values typical of long-distance dispersal. In Montcortès, C/H pollen was under the retting threshold during the whole Medieval epoch, suggesting that local/regional cultivation and long-distance dispersal were the main sources of this pollen. An abrupt increase took place at the beginning of Modern Age (1500-1600 CE) that exceeded the retting threshold and remained in this condition until late 19^th^ century, when a sharp decline occurred to long-distance dispersal values. During the last decades, a *C/H* pollen recovery was recorded compatible with regional cultivation, as no records of this activity exist around the lake.

**Figure 7.**
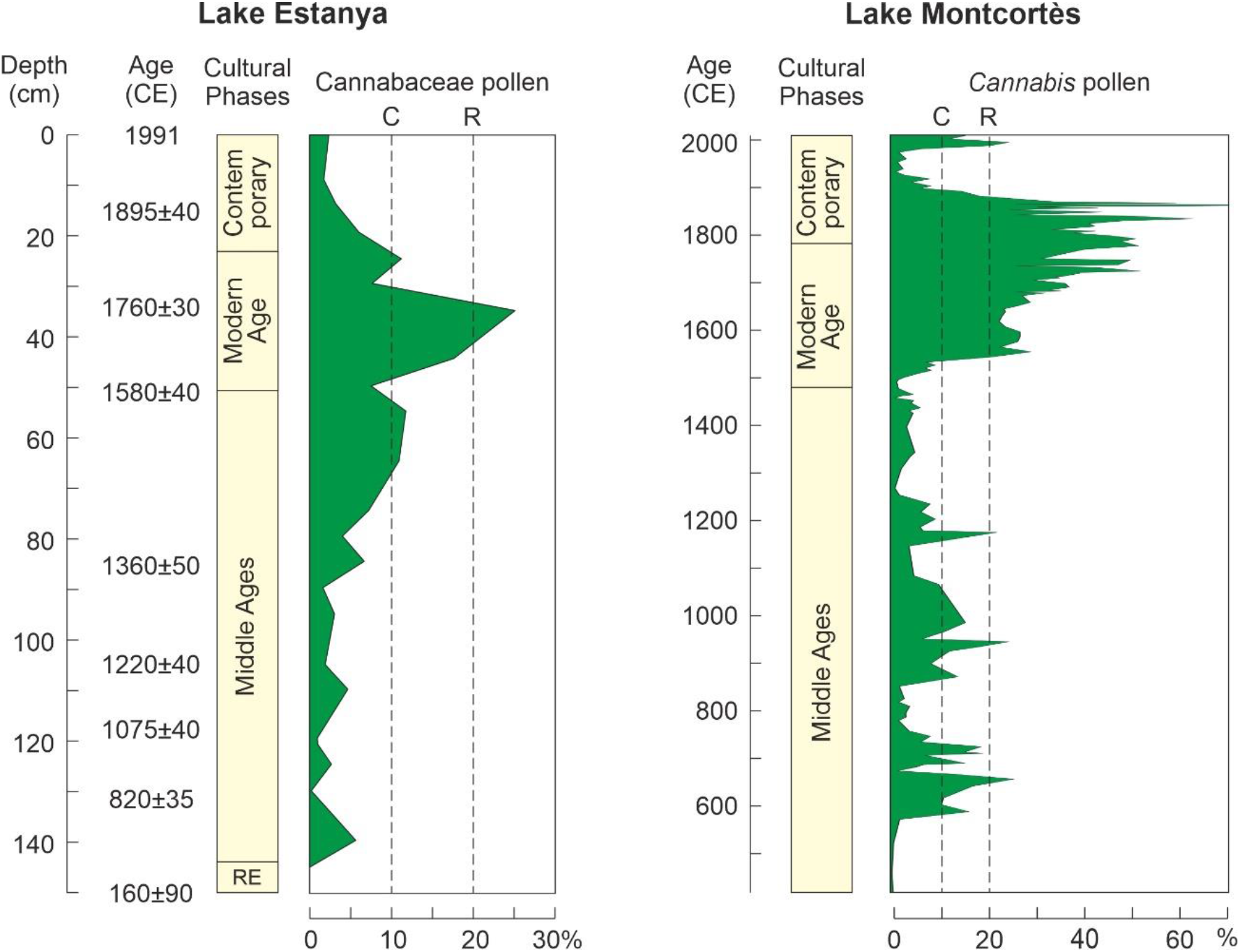
Comparison of the Estanya (Riera et al., 2004) and Montcortès (Rull et al., 2021b) *C/H* pollen records with indication of the thresholds suggested here for cultivation and hemp retting (C, cultivation threshold; R, retting threshold). The identification provided in the original references is given, although in this paper *C/H* pollen from these localities has been assigned to cultivated *Cannabis* using the assemblage approach (Table 2).

### 5.2. Pleistocene colonization

The issue of pre-anthropogenic arrival of *C/H* wild forms to the IP deserve special attention because the FAs recorded in this work for the IP are considerably older than those provided before for the Late Pleistocene arrivals to western Europe (Fig. 1). Dating reliability and correct taxonomic identification are essential to evaluate the first arrivals of *Cannabis* pollen to the IP. The oldest records (Middle Paleolithic) found correspond to the Gran Dolina (García-Antón, 1989), Bolomor (Ochando et al., 2019) and Teixoneres (Ochando et al., 2020a) sites, where *C/H* pollen was identified as “Cannabiaceae” or “Cannabinaceae”, which are unusual terms not recognized as formal taxonomic units but are here considered as indicative of *C/H* pollen type (Table 3). Using the assemblage approach, *C/H* pollen has been assigned to wild *Cannabis* in Gran Dolina, where Mediterranean scrubs dominated the landscape, but the available dating (Middle Pleistocene) is not precise enough to resolve the age of the involved sediments. The other two localities have more dating resolution but less taxonomic accuracy, as the dominance of mixed Mediterranean-mesophyllous forests prevents to unequivocally assign the *C/H* pollen to either wild *Cannabis* or *Humulus* using the assemblage approach

**Table 3.** Paleolithic/Pleistocene records of *C/H* pollen in the IP, indicating the main features related with dating, taxonomical identification and vegetation types, according to the original references. AA (assemblage approach) assignments according to McPartland et al. (2018).

| Site | First appearance | Dating | *C/H* pollen | Vegetation | AA assignment |
| --- | --- | --- | --- | --- | --- |
| GE | 11.4-10.9 ky BP | Radiocarbon | *Cannabis/Humulus* | Mixed Atlantic forest | Wild *Humulus* |
| SB | 13.4 ky BP | ND | Cannabaceae | ND | ND |
| LR | 14.5-10.5 ky BP | Radiocarbon | *Humulus/Cannabis* | Mixed Atlantic forest | Wild *Humulus* |
| AS | 24.8-23.2 ky BP | Radiocarbon | Cannabinaceae | Mixed Mediterranean forest | *C/H* |
| BA1 | 31 ky BP | U-series | *Cannabis* | Cold (glacial) steppe | Wild *Cannabis* |
| TO | 43.1 ky BP (MIS 3) | Radiocarbon | Cannabaceae | Mixed Mediterranean forest | *C/H* |
| TE | 100 ky BP (MIS 5c) | U-series | Cannabinaceae | Mixed Mediterranean forest | *C/H* |
| BO | 150 ky BP (MIS 5e) | Faunal | Cannabinaceae | Mixed Mediterranean forest | *C/H* |
| GD | Mid-Pleistocene | Faunal | Cannabiaceae | Mediterranean scrub | Wild *Cannabis* |

Therefore, the oldest well-dated occurrences of *C/H* pollen recorded in the IP correspond to the Middle Paleolithic of Bolomor and Teixoneres sites, ranging from 150 to 100 kyr BP (Mid-Late Pleistocene). These dates are similar to those previously recorded for eastern Europe and much older than those reported for central and western Europe (Fig. 1). This suggests that a first and previously undetected colonization wave would have reached the IP during the Mid-Late Pleistocene. The geographical origin of this first colonization wave is unknown but the fact that both localities involved are located on Mediterranean coasts suggests an east-west dispersal direction across this sea (Fig. 8). This Mediterranean pathway (MP) would require aerial and/or aquatic dispersal and would be favored by the existence of several islands and archipelagos (Fig. 1) favoring stepping-stone dispersal (Saura et al., 2014). MP dispersal would have occurred by animal-mediated terrestrial or aerial transport but eventual unintentional dispersal by human migrants cannot be ruled out (Clarke & Merlin, 2013).

**Figure 8.**
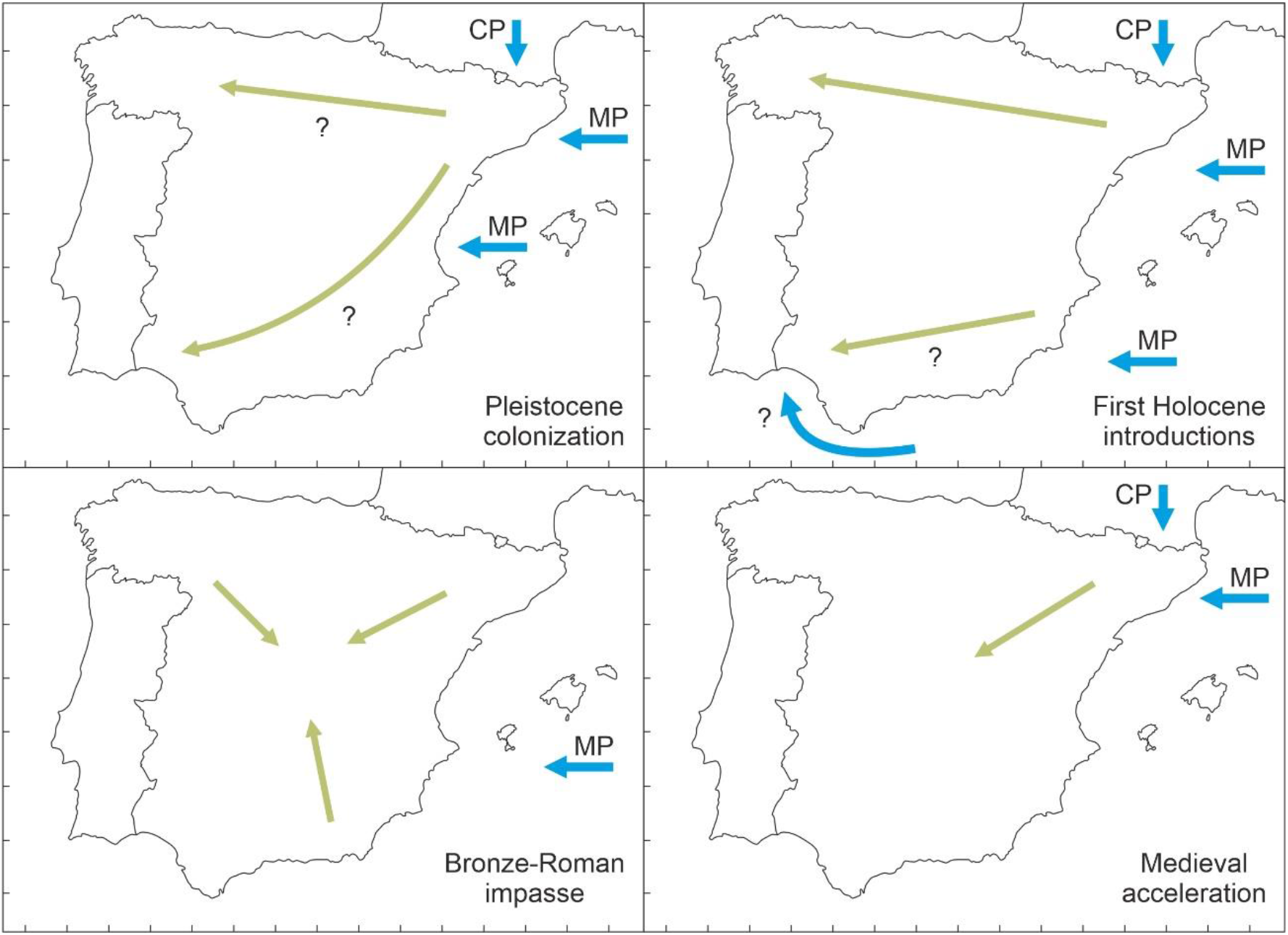
Possible dispersal pathways for *C/H* pollen into (blue arrows) and within (green arrows) the Iberian Peninsula.

The other Pleistocene FAs recorded in this study correspond to the Upper Paleolithic and range from ^~^43 kyr to ^~^13 kyr BP (Late Pleistocene). In this case, older FAs (^~^43-25 kyr BP) are situated in the NE and SW parts while the younger arrivals (^~^15-11 kyr BP) are in N and NW sectors, which could indicate the occurrence of two different colonization waves from outside before and after ^~^15-25 kyr BP or internal IP migration trends. The two older records are in the NW sector, which suggests the continued activity of the MP, the rise of a new continental pathway (CP), or both. This would support the idea of internal IP migrations after the colonization of NW coasts but the available evidence is insufficient for a sound appraisal. In a continental-wide perspective, the period between 18.5 kyr BP and 10.5 kyr BP was characterized in Europe by cold steppes in which wild *Cannabis* was the main *C/H* component, whereas *Humulus*-bearing deciduous forests began to spread from the southern glacial refugia (McPartland et al., 2018). In our study, a Late-Pleistocene record from the Mediterranean biogeographical region of the IP has been assigned to wild *Cannabis*, while two records from the Eurosiberian region have been considered to be consistent with wild *Humulus* (Table 3). No evidence for cultivated *Cannabis* has been found.

### 5.3. Holocene expansion/anthropogenic diffusion

Before discussing Holocene *C/H* pollen trends, it is necessary to introduce the chronological framework of ecological and landscape dynamics for SW Europe. Current European landscapes have been shaped during the last 10-12 millennia under the influence of climatic shifts and human impact, along with their corresponding feedbacks and synergies. The role of natural and anthropogenic drivers in the origin of Eurosiberian and Mediterranean biomes in SW Europe has largely been discussed but the increasing anthropization of the landscapes they conform is widely recognized, especially since the Neolithic (Bocquet-Appel et al., 2009, 2012; Collins et al., 2012; Leppard, 2014; Roberts et al., 2018; Vannière et al., 2016). Superimposed to this progressive anthropization, there is a climatic trend toward aridification that has also influenced the emergence and expansion of the Mediterranean biome across the whole circum-Mediterranean region, including the IP (Jalut et al., 2009; Carrión et al., 2010; Pérez-Obiol et al., 2011; Vannière et al., 2011; Ramos-Román et al., 2018). This section begins with a summary of the main Holocene anthropogenic and paleoclimatic trends, to provide the appropriate socio-environmental chronological context for the discussion of the most significant *C/H* pollen events – namely the Neolithic wave, the Bronze-Roman impasse and the Medieval acceleration.

#### 5.3.1. Anthropization chronology

The latest published chronological framework of human impact for SW Europe (Iglesias et al., 2021) subdivide the Holocene into four main stages, according to fire incidence, deforestation and land-use trends (Fig. 9):

- <u>Stage I</u> corresponds to pre-Neolithic times (>6.5 kyr BP) and was characterized by hunter-gatherer societies living in densely forested landscapes under climates with long, warm/dry summers and high interannual precipitation variability, which would have favored the occurrence of frequent fires.
- <u>Stage II</u> roughly coincides with the Neolithic (6.5-4.2 kyr BP) and was characterized by extensive deforestation and land use under cooler and wetter climates. During this stage, human impact increased and the SW European landscapes became more cultural and characterized by more open communities with frequent forest/cropland mosaics. Human societies were increasingly sedentary and fast-growing, and practiced a diversified economy.
- <u>Stage III</u> corresponds to the first half of the Late Holocene (4.2-2.4 kyr BP) and was characterized by high spatial climatic variability and the expansion of anthroposystems leading to substantial deforestation and the dominance of open landscapes, notably for cultivation and grazing. This was the time of decoupling of ecosystem dynamics from climate and human disturbance became the major ecological forcing. Human footprint on the landscape was especially important between 3.2 and 2.4 kyr BP (Bronze-Iron ages), which was the onset of more intense and efficient agriculture practices.
- <u>Stage IV</u> encompasses roughly the last two millennia (2.4 kyr BP to present) and was characterized by maximum rates of human impact, as expressed in intensive land use and the development of urban extractive industries. Burning and deforestation increased and peaked during the Roman Period (^~^2 kyr BP), possibly due to the need of expanding arable lands, and declined afterwards, when technological innovations facilitated more intensive exploitation practices. Large changes in transport and communication facilitated exchange and commerce after the collapse of the Roman Empire.

**Figure 9.**
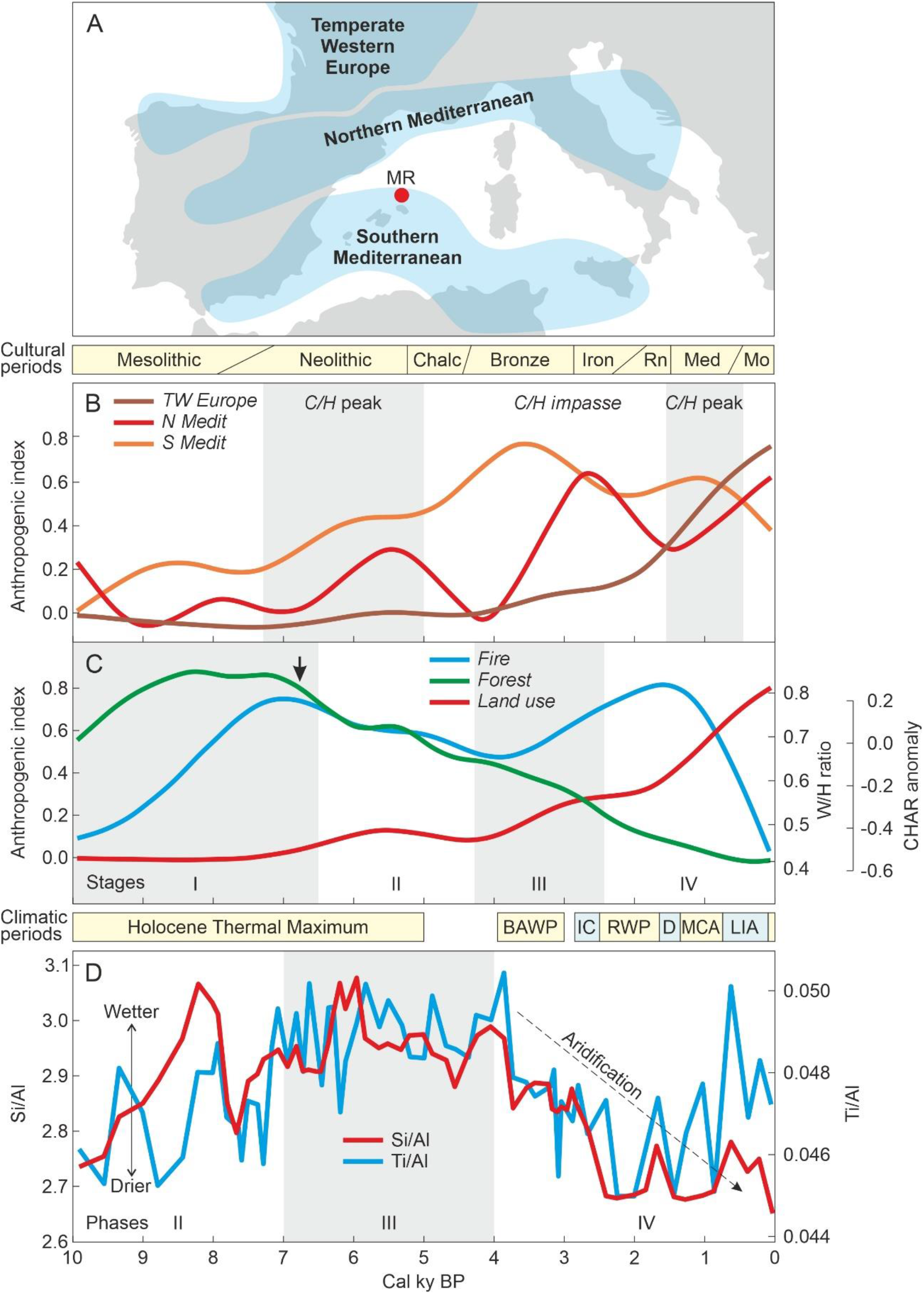
Holocene anthropogenic (B, C) and paleoclimatic (D) chronological frameworks for SW Europe. A) Sketch-map indicating the cultural subregions defined by Iglesias et al. (2021) and the location of the Minorca paleoclimatic record (red dot) (Frigola et al., 2007). B) Subregional differences in human impact as expressed by land-use values, using an anthropogenic index based on pollen percentages of cultivated plants (*Avena, Hordeum, Secale, Triticum, Zea*, other cereals, *Castanea, Juglans* and *Olea*) (Iglesias et al., 2021). The gray bands highlight the times of maximum *C/H* first appearances (see Figs. 2 and 6). C) Fire, forest cover and land use intensity for the whole SW European region. Fire incidence is expressed as anomalies in charcoal (CHAR) accumulation rates. Forest cover is expressed as the ratio between woody (W) and herbaceous (H) taxa, with high values representing closed forests and low values open vegetation. Land-use intensity is expressed by the previously defined anthropogenic index. The black arrow is the onset of pronounced forest lost (deforestation). I to IV are the stages of development of socio-environmental systems (Iglesias et al., 2021). D) Moisture trends reconstructed from the Minorca record (red dot) using the Ti/Al and Si/Al ratios from core MD99-2343. II to IV are the climatic phases defined on the basis of these ratios (Frigola et al., 2007). Cultural periods (Iglesias et al., 2021): Chalc, Chalcolithic; Rn, Roman Epoch; Med, Medieval Period; Mo, Modern Age. Climatic periods (Renssen et al., 2009; Martín-Chivelet et al., 2011): BAWP, Bronze Age Warm Period; IC, Ice Age Cold Period; RWP, Roman Warm Period; D, Dark Ages Cold Period; MCA, Medieval Climate Anomaly; LIA, Little Ice Age.

These stages should be viewed as a regional expression with subregional differences, especially in the chronological boundaries and the intensity of human impact (Fig. 9). For example, Stage II began roughly a millennium later (^~^5.1 kyr BP) in Temperate Western Europe (TWE), where Neolithization occurred later than in the Mediterranean region. Regarding human impact, as expressed in land-use intensity, the TWE trend is more similar to the monotonically increasing regional trend, whereas Northern and Southern Mediterranean regions show significant differences. In the north, the ascending trend was spiked by two maxima at ^~^5.5 kyr BP (Neolithic) and ^~^2.5 kyr BP (Iron Age), whereas in the south, the trend was increasing until a maximum at ^~^3.5 kyr BP (Bronze Age) and then decreased until Modern times (Iglesias et al., 2021).

#### 5.3.2. Paleoclimatic framework

Paleoclimatic trends for the western Mediterranean region have been characterized using a marine sedimentary record retrieved in the Balearic archipelago, the Minorca MD99-2343 record, which contains the whole Holocene sequence (Frigola et al., 2007). This record distinguished four main climatic phases, three of them occurring during the last ^~^10 ky BP (Fig. 9):

- <u>Phase II</u> (10.5-7 kyr BP), roughly coinciding with anthropogenic Stage I, characterized by increasing Si/Al and Ti/Al ratios, which are proxies for terrigenous input from fluvial sources and, therefore, for increasingly wetter continental climates.
- <u>Phase III</u> (7-4 kyr BP), roughly coinciding with anthropogenic Stage II, characterized by high and less variable terrigenous input, indicating wetter and fairly stable continental climates.
- <u>Phase IV</u> (4-0 kyr BP), encompassing anthropogenic stages III and IV, characterized by a maintained decrease of terrigenous input, consistent with a Late Holocene aridification trend.

Paleotemperature trends in the IP have followed the usual Northern Hemisphere sequence beginning with the Early Holocene Warming (up to ^~^11 kyr BP) followed by the Holocene Thermal Maximum (HTM; 11-5 kyr BP) (Renssen et al., 2009) and the higher frequency Late Holocene variability characterized by the Bronze Age Warm Period (4-3 ky BP), the Iron Age Cold Period (2.9-2.5 kyr BP), the Roman Warm Period (2.5-1.7 kyr BP), the Dark Ages Cold Period (1.7-1.4 kyr BP), the Medieval Climate Anomaly (1.4-0.8 kyr BP), the Little Ice Age (0.8-0.1 kyr BP) and the Modern Warming (Martín-Chivelet et al., 2011).

#### 5.3.3. The Neolithic wave

The first acceleration of *C/H* arrivals to the IP occurred during the Neolithic but there is a consistent spatiotemporal pattern extending from the Mesolithic to the Chalcolithic. The Mesolithic FA patterns support the predominance of the MP with no internal migrations within the IP. The occurrence of a non-Mediterranean SW locality would be compatible with the extension of the MP to the Atlantic after crossing the Strait of Gibraltar or the dispersal through the southern coasts (Fig. 8). The MP continued to be active during the Neolithic, especially in the NE sector, from where a pathway of internal colonization progressed toward the SW of the IP. Once more, the possibility of the combined action of a MP and a CP is strongly suggested and is reinforced by the arrangement of the FAs during the Chalcolithic. The NW Atlantic focus initiated in the Mesolithic remained active until this cultural period. These consistent patterns suggest that the first Holocene introductions of *C/H* into the IP would have been a continuous process initiated in the Mesolithic, approximately 2000 years after the end of Pleistocene colonization, and continued until the Chalcolithic, with an intermediate Neolithic acceleration.

Whether these introductions were natural or anthropogenic remains unclear. The assignment of several IP records to cultivated *Cannabis* (Table 2) seem to favor Neolithic cultivation but this would be contradictory with the general continental patterns showing no *Cannabis* cultivation during the Neolithic. Indeed, across Europe, the time interval between 10 kyr BP and 5 kyr BP was characterized by the appearance of the first crops and the expansion of deciduous mesophytic forests, which began to recolonize the continent since 7 kyr BP onward. In these conditions, most *C/H* pollen records were consistent with *Humulus*, whereas *Cannabis-prone* records were limited to the southern Mediterranean coasts. Interestingly, no pollen records consistent with cultivated *Cannabis* were found in the IP, which coincided with the lack of archaeological evidence on *Cannabis* cultivation and use during the European Neolithic (McPartland & Hegman, 2018). The first paleoecological and archaeological signs of European hemp cultivation have been found in the western coasts of the Caspian Sea (present-day Bulgaria) during the Copper (Chalcolithic) and Bronze ages (Fig. 1). The involved domestication process has been considered autochthonous and independent from the early Neolithic domestication of *Cannabis* in the present China, suggesting that hemp domestication could have occurred independently in more than a single region (McPartland et al., 2018).

The beginning of the Neolithic *C/H* peak coincided with the onset of the general deforestation in SW Europe and occurred mainly during anthropogenic Stage II, when land use began to intensify (Fig. 9). During this stage, human impact underwent a significant increase, especially in the Mediterranean region, which is consistent with a higher CP (northern Mediterranean subregion) and MP (southern Mediterranean subregion) activity. Climatically, the Neolithic *C/H* wave coincided with a cooling trend at the end of the HTM and the stabilization of wetter climates in the western Mediterranean region, coinciding with the end of the postglacial sea-level rise to present-like settings (Frigola et al., 2007). This would have favored navigation and, hence, the maritime MP for *C/H* arrivals to the IP would have been especially favored. The EP should also be considered as a potentially important incoming pathway, as Indo-European peoples were important colonizers of the IP during this prehistorical phase and used preferably the Pyrenean pathway, situated to the north.

#### 5.3.4. The Bronze-Roman impasse

*C/H* introductions to the IP significantly declined between the Bronze Age and the Roman period, when a centripetal diffusion toward the central part of the IP began (Fig. 8). The few arrivals recorded from outside occurred in the SE and the Balearic Islands, likely following the MP. This can be regarded as a phase with little activity in relation to *C/H* expansion, when the cessation of the EP and the initial colonization of the most interior lands of the peninsula would be the most relevant features. Curiously, the Bronze-Roman impasse observed in the IP occurred shortly after the purported European domestication of hemp and during the expansion of this crop across the continent, which began by 4.5 kyr BP (Fig. 1) and intensified since 2 kyr BP, with the expansion of the Roman Empire (McPartland et al., 2018).

The Bronze-Roman impasse roughly coincided with anthropogenic Stage II, when deforestation of SW Europe was in progress and biomass burning experienced a relevant increase coinciding with a second regional pulse of human impact (Fig. 9). This enhanced anthropogenic pressure was especially significant in the Mediterranean region, peaking at the Bronze Age (southern subregion) and the Bronze-Iron transition (northern subregion). Contrastingly, maximum rates of regional land use coincided with a slowdown in *C/H* arrivals to the IP, where centripetal diffusion was the predominant process. This was an epoch of frequent arrivals to the IP, including peoples such as the Phoenicians, Greeks, Carthaginians or Romans, with well-developed navigation skills. Therefore, the low rates of *C/H* input cannot be attributed to the lack of new colonizers. It is possible that these incoming peoples were less attached to the hemp industry that former Neolithic colonizers or that they used the already existing local hemp resources with no need for new introductions. Archaeological evidence is needed to test this possibility and others that may emerge (Guerra & López-Sáez, 2006).

#### 5.3.5. The Medieval acceleration

A completely different pattern emerged in the Middle Ages with the onset of the second diffusion wave. During this phase, older arrivals where concentrated in the NE sector, suggesting the reactivation of the northern CP, combined or not with the MP, that time restricted to the NE area (Fig. 8). From there, *C/H* diffusion advanced toward the central IP, which was colonized by *C/H* mostly during the Medieval and Modern periods. This phase falls within the European expansion of cultivated *Cannabis* recorded between 2 and 0.8 kyr BP (McPartland et al., 2018).

The significant increase of *C/H* pollen recorded in some localities (Estanya, Montcortès) during Modern Ages has been associated to hemp retting (section 5.1) and coincided with a phase of maximum development of the Spanish royal navy after the Columbian discovery of America (1492 CE) (Riera et al., 2006; Rull & Vegas-Vilarrúbia, 2014; Trapote et al., 2018). During that phase, hemp cultivation was mandatory in the whole country to provide fiber, mainly for sails and ropes, to support the naval expansion of the Spanish Empire (Sanz, 1995). The maximum of hemp industry recorded during the mid-18^th^ century coincided with a phase of maximum ship building, especially in the NE part of the IP, and general economic prosperity and intensification of commerce with America (Andreu, 1981; Delgado, 1994). Hemp industry was severely curtailed in the mid-19^th^ century after the dismantling of the royal navy and the replacement of hemp fiber by other products, notably flax, since 1860 CE (Casassas, 1985).

The Medieval acceleration coincided with a third pulse in regional human impact and the continued general deforestation, along with a significant decline of biomass burning that has been attributed to the development of new methods of resource management more efficient that fire (Fig. 9). Human pressure was growing faster in temperate western Europe and the northern Mediterranean region, which is consistent with the *C/H* arrival and expansion from the peninsular NE (Fig. 8). The pattern is also consistent with the progressive colonization of the IP from the north by Christian cultures (with important contributions from beyond the Pyrenees) during Medieval times, after the southern Muslim invasion (Brunes Ibarra, 2006; Blasco & Badia-Miró, 2007; Alonso et al., 2022).

### 5.4. Possible influence of climatic changes

Finding direct evidence for a causal link between climatic shifts and *C/H* expansion across the IP is difficult but eventual chronological coincidences may suggest some potential relationships. The oldest (Middle Paleolithic) well dated *C/H* records of the IP – i.e., Bolomor (150 kyr BP), Teixoneres (100 kyr BP) and Toll (^~^43 kyr BP) – appeared during interstadials MIS 5e, MIS 5c and MIS 3, respectively (Ochando et al., 2019, 2020a, b), which may suggest some link with warmer climates. The Asperillo record (^~^25-23 kyr BP) occurred in a short warming between the cooling reversals known as Heinrich Event 2 (HE2) and the Last Glacial Maximum (LGM) (Fernández et al., 2021), which may support a potential relationship with temperature. Upper Paleolithic records (La Roya, Son Bou and Gesaleta) occurred during the Lateglacial transition (^~^15-12 kyr BP), with no clear links with any specific climatic conditions.

Regarding Holocene times, the Neolithic colonization wave (7.2-5 kyr BP) coincided with the global HTM (Renssen et al., 2012), whereas the Bronze-Roman impasse (^~^4-2 kyr BP) occurred under variable climatic conditions in the IP (Martín-Chivelet et al., 2013). The Medieval diffusion wave (1.5-0.5 kyr BP) took place mostly during the North Atlantic warming known as the Medieval Warm Period (MWP) or the Medieval Climate Anomaly (MCA) (Mann et al., 2009). As it occurred in the Pleistocene records, the coincidence between *C/H* appearances/bursts and warmer climates seems to be fairly consistent. It is also noteworthy that this relationship does not maintain in cases where the *C/H* pollen has been assigned to *Humulus*, rather than *Cannabis*, by the assemblage approach, as it occurs in the Lateglacial (Table 3). Therefore, the potential link between *C/H* pollen and warmer climates seems to be more consistent for *Cannabis*, which is consistent with the general climatic requirements of this plant (Clarke & Merlin, 2013). In contrast with Pleistocene times, the influence of climate during the Holocene could have been indirect by affecting human migration and settlement patterns, with the ensuing consequences on *C/H* diffusion.

Relationships with moisture changes are more difficult to establish, as shifts in hydrological balance, i.e. precipitation/evapotranspiration rates, are usually of more local nature than temperature variations. In this case, the establishment of local pollen-climate relationships is recommended, followed by the seek of more general regularities across spatial scales. However, the Late-Holocene aridification trend was a regional phenomenon whose potential influence on *C/H* pollen trends may be interesting to evaluate, as it has been considered a major environmental driver for the origin and expansion of Mediterranean biomes. Noteworthy, two contrasting *C/H* events – i.e., the Bronze impasse and the Medieval acceleration – took place during this trend toward drier climates. In addition, the Neolithic *C/H* acceleration occurred during a phase of wet climates, which differs from the aridity trend under which the same phenomenon developed during the Middle Ages. Therefore, a potential causal relationship between aridity and *C/H* dispersal is not supported by these preliminary results. In the absence of more detailed studies, it seems that temperature would be more influential than moisture on *C/H* biogeography in the IP, but more climatically-focused research is needed for a sound assessment.

## 6. Future research

As this is the first time that a synthetic reconstruction of spatiotemporal patterns of *C/H* pollen across the IP is attempted, the results obtained should not be considered conclusive assessments but, rather, a source for hypothesis and ideas to be tested with future studies. As usual in any meta-analysis, the validity of the outputs will expire when a new similar synthetic study will be produced using an updated database.

As pointed out in the introduction, the database used here has been extracted from the exhaustive compilation of palynological records available for the IP (Carrión et al., 2012, 2022), enriched with personal contributions from the authors of a number of records that remain to be published or whose *C/H* pollen records are not available in the original publications. Further efforts should be aimed at increasing and improving the *C/H* records of this database, to pave the way towards a new and more exhaustive meta-analysis. This is a long-term venture worth to be continued because of the many implications that *Cannabis* – as a most ancient human domesticate, with a number and a diversity of uses difficult to match (Balant et al., 2021) – may have for the study of cultural evolution, in relation to the domestication and use of natural resources.

The IP is especially well suited for a study of this type due to its strategic geographical emplacement, which is responsible not only for its outstanding environmental and biogeographical patterns but also for its rich cultural history, in terms of both incoming and outgoing influence. Indeed, the IP has been the meeting place of a high diversity of cultures since prehistoric times but has also influenced global cultural developments of the modern world after the Columbian discovery of America and the further worldwide imperial expansion. As shown in this paper, most of these processes can be linked, in one or another way, to the historical biogeography of the Iberian *Cannabis*.

As emphasized in the introduction, this study should be complemented with a similar peninsular-wide meta-analysis of archaeological and historical evidence, to establish the corresponding links of *Cannabis* arrival, dispersal and diffusion, as recorded by pollen evidence, with human migrations and cultural changes across spatial and temporal scales.

## Acknowledgements

No funding was received specifically for the development of this research.

## Notes

### Competing Interest Statement

The authors have declared no competing interest.

### Summary of Updates

This is the same version submitted before (nothing has changed in the pdf) but co-authors, who were missing in the online form of the former version, have been added

